# Recovery mode: Marine protected areas enhance the resilience of kelp species from marine heatwaves

**DOI:** 10.1101/2024.05.08.592820

**Authors:** Carolina Olguín-Jacobson, Nur Arafeh-Dalmau, Michelle-María Early-Capistrán, José Antonio Espinoza Montes, Arturo Hernández-Velasco, Ramón Martínez, Alfonso Romero, Jorge Torre, C. Brock Woodson, Fiorenza Micheli

## Abstract

Marine protected areas (MPAs) can promote population recovery from climate change impacts by reducing local stressors, such as fishing. However, with extreme climatic events such as marine heatwaves (MHWs) increasing in frequency and duration, it remains unclear whether MPAs enhance recovery following these acute perturbations, and how recovery varies across species and ecological traits (e.g., sedentary vs mobile species). We used 16 years (2007-2022) of kelp forest monitoring data in Isla Natividad, Baja California Sur, Mexico, to assess the impact of the 2014-2016 MHWs on fish and invertebrate communities. Then we evaluated the impact and recovery from the MHWs of economically and ecologically important invertebrate species inside and outside two fully protected marine reserves. We found that the 2014-2016 MHWs, which were the most intense and persistent ever observed in Isla Natividad, impacted invertebrates but not fish biomass. Marine reserves did not confer resistance to the MHWs, however, reserves did enhance the recovery of some species after the MHWs. Inside marine reserves, abalone (*Haliotis* spp.) and wavy turban snail (*Megastraea* spp.) (benthic sedentary invertebrates) recovered to pre-heatwave biomass after two years and spiny lobster (*Panulirus interruptus*) (benthic mobile invertebrate) after four years. Outside the reserves, abalone recovered after three years, while the other two species never recovered. The warty sea cucumber (*Apostichopus parvimensis*) population collapsed after the MHWs and never recovered inside nor outside the reserve. Remarkably, abalone biomass had an outstanding and sustained recovery inside reserves, with a 5.6-fold increase in biomass after the MHWs, which was over three times higher than the recovery reported outside the reserve. Our analysis of long-term monitoring data shows that marine reserves cannot prevent adverse impacts from extreme climatic events but can enhance species recovery following these events. Benefits conferred by marine reserves, however, are species-specific and may be limited to species with limited dispersal and localized population dynamics.

## 1. Introduction

Marine protected areas (MPAs) promote biodiversity and ecosystem functioning and are the cornerstone of many conservation and management strategies (Grorud-Colvert *et al*. 2021). MPAs contribute to the recovery of overexploited species (Edgar *et al*. 2014; Sala *et al*. 2018) and can enhance the resilience and adaptive capacity of species and ecosystems to climate impacts (Micheli *et al*. 2012; Roberts *et al*. 2017; Jacquemont *et al*. 2022; Benedetti-Cecchi *et al*. 2024). As a result, MPAs have been implemented globally to protect biodiversity, manage fisheries, and support the provision of ecosystem services, including tourism, recreation, food provision and carbon sequestration (Aburto-Oropeza *et al*. 2011; Sala *et al*. 2013; Gill *et al*. 2017). However, despite decades of research on the ecological and socioeconomic benefits of MPAs (Lester *et al*. 2009; Bates *et al*. 2014; Edgar *et al*. 2014), the benefits that MPAs can confer under climate change impacts remain less understood (Bates *et al*. 2019; Bruno *et al*. 2019).

Well managed and fully protected (no-take) MPAs have been shown to be most effective for recovering and maintaining the biomass of overfished species (Edgar *et al*. 2014), preserving genetic diversity (Munguía-Vega *et al*. 2015), enhancing ecosystem recovery following disturbance (Eisaguirre *et al*. 2020) and providing resilience against climate change (Micheli *et al*. 2012; Roberts *et al*. 2017; Ziegler *et al*. 2023; Benedetti-Cecchi *et al*. 2024). A notable example of a successful no-take marine reserve is the case of Cabo Pulmo National Park in the Gulf of California, Mexico, where, after 14 years of implementation, there was an increase of 463% of total fish biomass and 30% annual increase in predatory fish (Aburto-Oropeza *et al*. 2011). Recent global reviews and meta-analyses (Roberts *et al*. 2017; Jacquemont *et al*. 2022) found that no-take MPAs significantly benefit ecosystems and ecosystem services by increasing carbon sequestration, coastal protection, biodiversity, and the size and reproductive capacity of organisms, as well as help mitigate and promote adaptation to climate change.

Climate change is increasing the frequency, intensity, and duration of extreme climatic events such as marine heatwaves (MHWs; (Oliver *et al*. 2019; IPCC 2023)) that are impacting marine ecosystems worldwide (Smale *et al*. 2019; Wernberg 2021; Garrabou *et al*. 2022). As nations have committed to protect 30% of the oceans by 2030 while adapting to climate change (CBD 2022), a key question is how MHWs impact the function of MPAs, and whether MPAs contribute to ecological resilience in the face of extreme climatic events. In 2014-2016, the Northeast Pacific Ocean was subject to the strongest and longest MHWs ever recorded (Laufkötter *et al*. 2020), that vastly impacted marine ecosystems along the California Current system, from Alaska, USA, to Baja California, Mexico (Cavole *et al*. 2016). The 2014-2016 MHWs resulted in shifts in the distribution of species (Sanford *et al*. 2019), changes in community biomass and structure (Arafeh-Dalmau *et al*. 2019; Jiménez-Quiroz *et al*. 2019; Beas-Luna *et al*. 2020), loss of habitat-forming species such as kelp forests (Cavanaugh *et al*. 2019; McPherson *et al*. 2021; Arafeh-Dalmau *et al*. 2023b) and decline in, and closure of fisheries in the region (Cheung *et al*. 2020; Free *et al*. 2023).

Kelp forests are one of the most productive ecosystems on Earth, supporting a vast diversity of marine life (Schiel *et al*. 2015; Wernberg *et al*. 2019) and providing ecosystem goods and services that coastal communities depend on, including valuable fisheries (Wernberg *et al*. 2019; Eger *et al*. 2023). Kelp forests within the California Current were significantly affected by the 2014-2016 MHWs, but the impacts and recovery varied across regions (Cavanaugh *et al*. 2019; Bell *et al*. 2023). For example, in northern California, a >90% bull kelp (*Nereocystis luetkeana*) forest loss led to the collapse of the red abalone (*Haliotis rufescens*) and red sea urchin (*Mesocentrotus franciscanus*) fisheries, and an ecosystem shift from kelp forest to sea urchin barrens (Rogers-Bennett *et al*. 2019). Notably, these bull kelp ecosystems have not recovered as of 2021 (Arafeh-Dalmau *et al*. 2023b). In Baja California, Mexico, near the southern distribution limit of giant kelp (*Macrocystis pyrifera*), there was a significant decline in giant kelp forests, ∼50% decrease of fish species richness, and almost complete depletion of sessile invertebrates (Arafeh-Dalmau *et al*. 2019).

Isla Natividad, off the coast of Baja California Sur, Mexico, has one of the most persistent and stable kelp forests in the California Current (Arafeh-Dalmau *et al*. 2021) despite being located near the southern distribution range limit of giant kelp. Such persistence is possibly due to a combination of low coastal population density and low human impacts, small-scale local oceanographic processes that provide climate refugia, and effective management by local fishers (Woodson *et al*. 2019; Smith *et al*. 2022; Micheli *et al*. 2024). A fishing cooperative on Isla Natividad, known as *Buzos y Pescadores de la Baja California*, relies on different kelp-associated species, such as abalone, lobster, turban snail and sea cucumber for which the cooperative holds a concession for exclusive access and is responsible for local management and enforcement of regulations (McCay *et al*. 2014). In 2006, the fishing cooperative voluntarily established two no-take marine reserves (Fig. 1a), considering biological and economic factors when selecting the location and size of the reserves and investing in local enforcement and monitoring (Micheli *et al*. 2012; Micheli *et al*. 2024).

**Figure 1.**
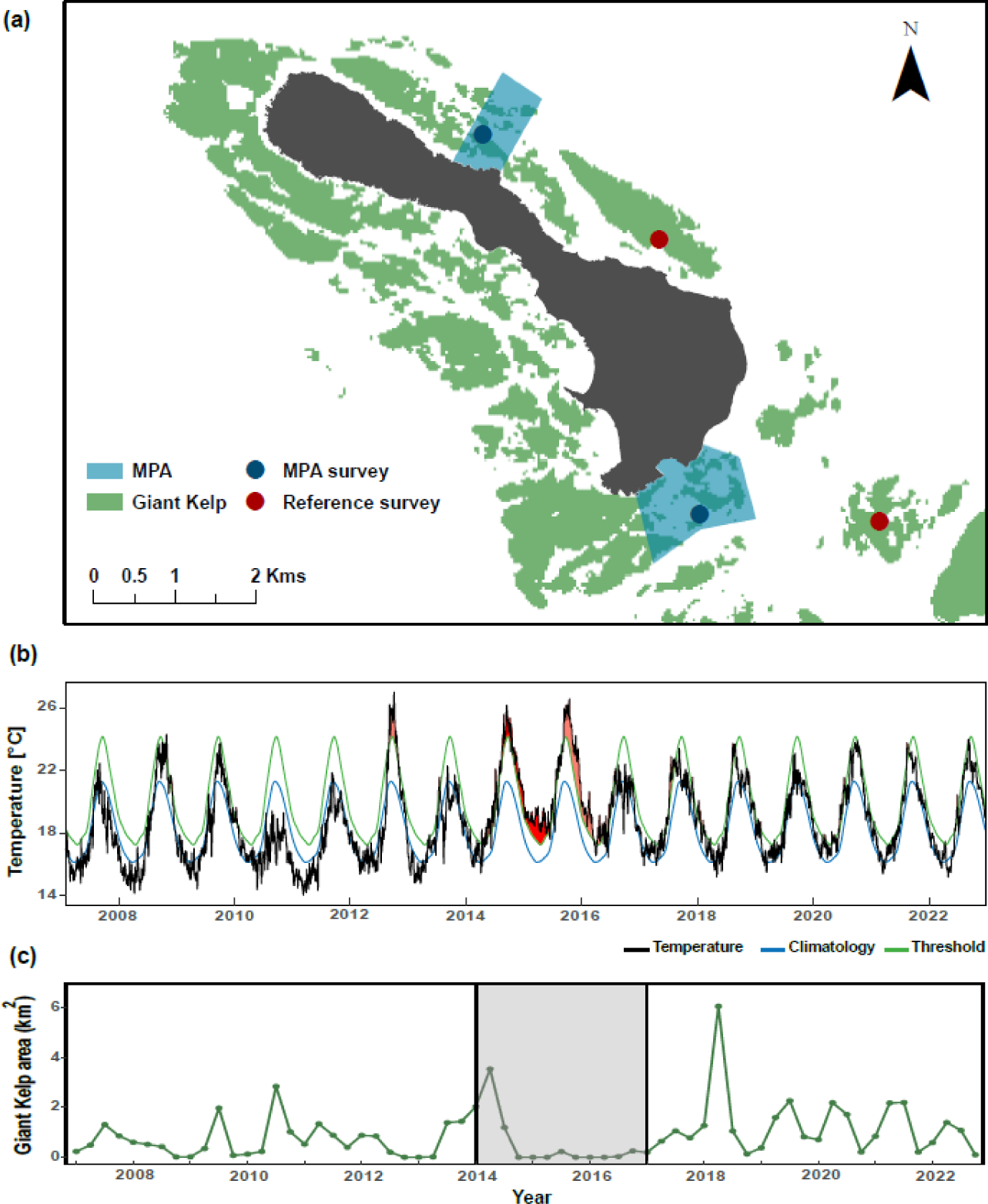
a) Map of the study area around Isla Natividad, showing the two marine reserve polygons and dots representing the reserves and reference survey sites (14-27 transects surveyed per year: dots represent the centroid around where surveys were conducted throughout the area). b) Time series of the sea surface temperature (SST) in Isla Natividad over the study period (2007-2022), with color lines indicating the MHWs (see Methods). c) Time series of the area (km^2^) of kelp canopy detected in each quarter of a year in Isla Natividad over the study period (2007-2022) (see Methods), gray shaded area represents the 2014-2016 MHWs.

Long-term monitoring is needed to quantify the possible benefits conferred by MPAs to species, populations, and ecosystems in the face of climate change impacts, given that the response and recovery of an ecosystem after a climatic event can span from months to years. Recovery is defined here as the rate of return to community structure and function similar to the pre-perturbation state (Capdevila *et al*. 2020). Previous studies have found that the marine reserves of Isla Natividad played a critical role in promoting the recovery of depleted abalone populations following mass mortality events (Micheli *et al*. 2012; Rossetto *et al*. 2015; Aalto *et al*. 2019; Smith *et al*. 2022). In 2009, prolonged hypoxia caused mass mortality of pink and green abalone (*Haliotis corrugata* and *H. fulgens*) populations. Population surveys and recruitment studies conducted during this event showed that reserves maintained higher abundances of larger reproductive individuals and greater juvenile recruitment within and around the reserves, highlighting the potential role of protection in supporting recovery by maintaining reproductive output and juvenile recruitment (Micheli *et al*. 2012). Continued monitoring confirmed that both species had a rapid recovery after a six-year fishing closure and the enforcement of the voluntary marine reserves. Pink abalone returned to the lower bound pre-mortality levels inside and outside the reserves, while green abalone recovered faster inside marine reserves (Smith *et al*. 2022). However, no studies have examined the generality of these benefits, and whether MPAs may promote the recovery of other invertebrate species after MHWs.

Here, we first characterize the 2014-2016 MHWs in Isla Natividad and describe the impact to giant kelp forests using remote sensed data. Then, we analyse 16 years (2007-2022) of kelp forest biological monitoring data to assess the responses of the fish and invertebrate community to the 2014-2016 MHWs. We then evaluated whether the 2014-2016 MHWs impacted the species biomass and whether they recovered after the event, inside and outside two fully protected marine reserves in Isla Natividad, Mexico. We hypothesized that 1) both groups were significantly impacted during the MHWs, 2) impacts were less severe inside marine reserves compared to reference sites, and 3) species recovered more rapidly (i.e., required fewer years to recover) to pre-heatwave species biomass inside marine reserves than at reference sites. To analyse the recovery, we chose four species of invertebrates (abalone (*Haliotis* spp.), the wavy turban snail (*Megastraea* spp.), the warty sea cucumber (*Apostichopus parvimensis*) and the California spiny lobster (*Panulirus interruptus*)). We did not include fish, because our analyses found that the fish community was not impacted by the MHWs. These species were selected because they were abundant before the MHWs and support important commercial export fisheries.

## 2. Materials and Methods

### 2.1 Study region

Isla Natividad is located on the Pacific coast of Baja California Sur, Mexico (Fig. 1a). The fishing cooperative *Buzos y Pescadores de la Baja California* on Isla Natividad holds exclusive fishing rights within defined fishing grounds (concession) around the island since 1942 (Crespo-Guerrero *et al*. 2018). In 2006, the fishing cooperative voluntarily established two no-take community marine reserves, (La Plana-Las Cuevas, 1.3km^2^ in surface area and Punta Prieta, 0.7km^2^) covering a total surface of 2km^2^. Since 2006, the cooperative has maintained a successful community-based management program that enforces no-take regulations and continuously monitors fishing and poaching activities around the island (McCay *et al*. 2014; Micheli *et al*. 2024). These reserves were officially established by the Mexican government as fish refuges from 2018 to 2023 (DOF 2018), before and after this period, reserves have been voluntary.

### 2.2 Marine heatwaves

We identified MHWs which are warm periods where temperature exceeds the 90^th^ percentile threshold relative to a baseline climatology (seasonal varying mean) and last for at least five consecutive days (Hobday *et al*. 2016). We obtained daily sea surface temperature (SST) estimates for Isla Natividad between 1982 and 2022 using the NOAA 0.25° grid-resolution OISST dataset (Huang *et al*. 2021). We then used the R package heatwaveR (Schlegel *et al*. 2018) to identify MHWs registered from 1982 to 2022 (Fig. 1b), relative to a 30-year baseline climatology for 1983 to 2012. We identified the start and end dates and calculated the duration and intensity for each MHW event.

### 2.3 Surface canopy biomass of giant kelp

We used an existing dataset that uses multispectral Landsat images to estimate the surface canopy area of giant kelp (*M. pyrifera*) forests (Bell *et al*. 2020) to characterize giant kelp dynamics around Isla Natividad. This dataset provides quarterly estimates of kelp canopy area at a 30 m grid resolution from 1984 to present. We aggregated all 30 m grid pixels that overlay with the fishing concession polygon of Isla Natividad and created a dataset from 2007 to 2022 (Fig 1c). This published dataset can be visualized on kelpwatch.org (Bell *et al*. 2023).

### 2.4 Ecological monitoring

Ecological monitoring was conducted every year between July-September from 2007 to 2022 within the two marine reserves and two reference sites (where fishing is allowed) with similar habitat characteristics to the marine reserves (Fig. 1a). Using SCUBA, trained divers lay a 30 x 2 m (60 m^2^) belt transect and record *in situ* abundance data of ecological and economically important fish and invertebrate species. Between 14-27 transects were surveyed at each site once/year, (av. = 19.93, SE = 0.40) at depths ranging from 4-25m. Then, we calculated the biomass of fish and invertebrate species based on mean biomass estimates obtained from Woodson *et al*. (2019) and from Fajardo-León *et al*. (2008) for warty sea cucumber (*A. parvimensis*) (Table S1).

### 2.5 Statistical Analyses

#### Replication statement

| Scale of inference | Scale at which the factor of interest is applied | Number of replicates at the appropriate scale |
| --- | --- | --- |
| Site type | Pooled sites (reserve and reference) | 2 reference sites<br>2 reserves sites |
| Site-specific biomass | Pooled sites (reserve and reference) | 1275 of each combination of species (*Haliotis* spp., *Megastrea* spp., *A. parvimensis*, *P. interruptus*) and site type (reference/reserve) |

We employed Generalized Linear Models (GLMs) with Gaussian link functions in R version 4.3.2 (Team 2019) to analyze our data. Initially, we focused on examining the response of two broad taxonomic groups, fish and invertebrates, assessing their biomass trends (kg/60 m^2^) from 2007 to 2016 in relation to the predictor variables protection status (reserve/reference), heatwave status (before/during), and year. Before conducting our analyses, we standardized the data by pooling it into reserve and reference site categories, accounting for spatial scale differences. Additionally, we standardized biomass to annual mean values to address sampling variability across sites and log-transformed biomass values while adding a constant of 1 to handle zero values. This enabled us to determine whether these species experienced significant impacts during MHWs and whether such impacts varied between marine reserves and reference sites.

We did not calculate the recovery for fish species because the modeling showed that they were not significantly impacted by the MHWs. Therefore, we only focused on invertebrate species since they were impacted by the MHWs. We applied the same data treatment process and predictor variables in further analyzing the recovery of the four most economically important invertebrate species at Isla Natividad: *Haliotis* spp.*, Megastraea* spp., *A. parvimensis, and P. interruptus*. We iteratively ran models for each combination of predictor variables, both with and without interactions, until achieving a parsimonious model with statistically significant effects (α = 0.05), substantial explanatory power assessed through Cox-Snell R^2^ and relatively low Akaike Information Criterion (AIC) values, and robust residuals (ei ∼ N(0, σ^2^)). For H*aliotis* spp., *Megastraea* spp. and *P. interruptus*, we employed Gaussian link functions and evaluated model assumptions via residual analysis through visual inspection and statistical tests, ensuring zero mean (t-test, p > 0.05), normal distribution (Shapiro-Wilke test, p > 0.05), homoscedasticity (Levene’s test, p > 0.05), and independence (Ljung-Box test, p > 0.05; (Inchausti 2022)). For *A. parvimensis*, for which biomass values ranged from 0 to 1, we compared Gaussian and quasibinomial link functions and assessed model fit through residual deviance analysis. This comprehensive approach ensured the robustness and reliability of our findings across all species.

#### Species response and recovery to the 2014-2016 MHWs

To complement the modeling approach, we assess the impacts and recovery of species biomass during and after the MHWs by evaluating the percentage change in mean annual biomass (henceforth “biomass change”) per transect (kg/60 m^2^). First, during the MHWs (2014-2016) compared to the mean baseline values before the MHWs (2007-2013) and then after the MHWs (2017-2022) compared to the mean values during the MHWs. To assess recovery in relation to pre-heatwave values, we compared the yearly percentage change after the MHWs to the baseline annual mean values before the MHWs. We defined an early recovery signal as the first year when the species abundance reached or exceeded the lower bound of pre-MHWs abundance, which we defined as the 25th percentile of baseline values.

## 3. Results

### 3.1 MHWs

Isla Natividad was subject to 677 MHW days between 2014-2016, representing 38% of all MHW days registered in the past four decades (1983-2022) (Fig. 1b, Table S2). The most prolonged MHW lasted 302 days, from August 13, 2014, to June 10, 2015. The year with the highest number of MHW days registered was 2015, with 308 days and with a yearly cumulative MHW intensity of 908.3 °C days, followed by 2014 (235 days and 595.7 °C days) and 2016 (153 days and 334.4 °C days) (Table S2 and S3). The maximum intensity registered during the 2014-2016 MHWs peaked at 5.7 °C in October 2015, with temperatures reaching 26.6 °C. However, this was not the highest maximum temperature registered during a MHW in the time series. In October 2012, a MHW that lasted for 62 days reached a maximum intensity peaking at 5.9 °C, with temperatures reaching 27 °C (Fig. 1b, Table S2). The 2014-2016 MHW event was the strongest and most intense registered in Isla Natividad, with yearly cumulative intensities being ∼2 times higher than previous strong El Niño events of 1983, 1992, and 1997 (Table S2 and S3).

### 3.2 Kelp Biomass

We qualitatively document the loss and recovery of kelp forests during the 2014-2016 MHWs at Isla Natividad (Fig. 1c). After spring 2014, the satellite did not detect kelp forest at the surface in Isla Natividad, and the area remained at zero or very low values until the end of 2016. In 2017, kelp forest recovered to pre-MHW coverage and remained that way until the end of this study. As expected, kelp area also dropped to zero values after summer 2012, coinciding with the strongest SST temperatures ever registered during a MHW, and remained negligible until spring 2013 when it recovered (Fig. 1b-c).

### 3.3 Impact of MHWs and marine reserves benefit to commercial species

Generalized Linear Models (GLM) revealed that the MHWs had significant negative impacts on the biomass of invertebrates but not on the biomass of fish (Table S4) (Fig. 2a & 2b). The greatest decrease in biomass during the MHWs (2014-2016) for both kelp related groups and all invertebrate species was recorded in 2016, both inside and outside the reserve (Fig. 2a-f). The biomass of all species declined between 40-95% (Fig. 3a) during the MHWs compared to the pre-heatwave period. However, the decline was less severe inside the reserves than in the reference sites for all species (57-94% loss in reference sites and 39-86% loss in reserves) except for abalone that had a bigger decline in reserves (82%) than in reference (71%) sites (Fig. 3a).

**Figure 2.**
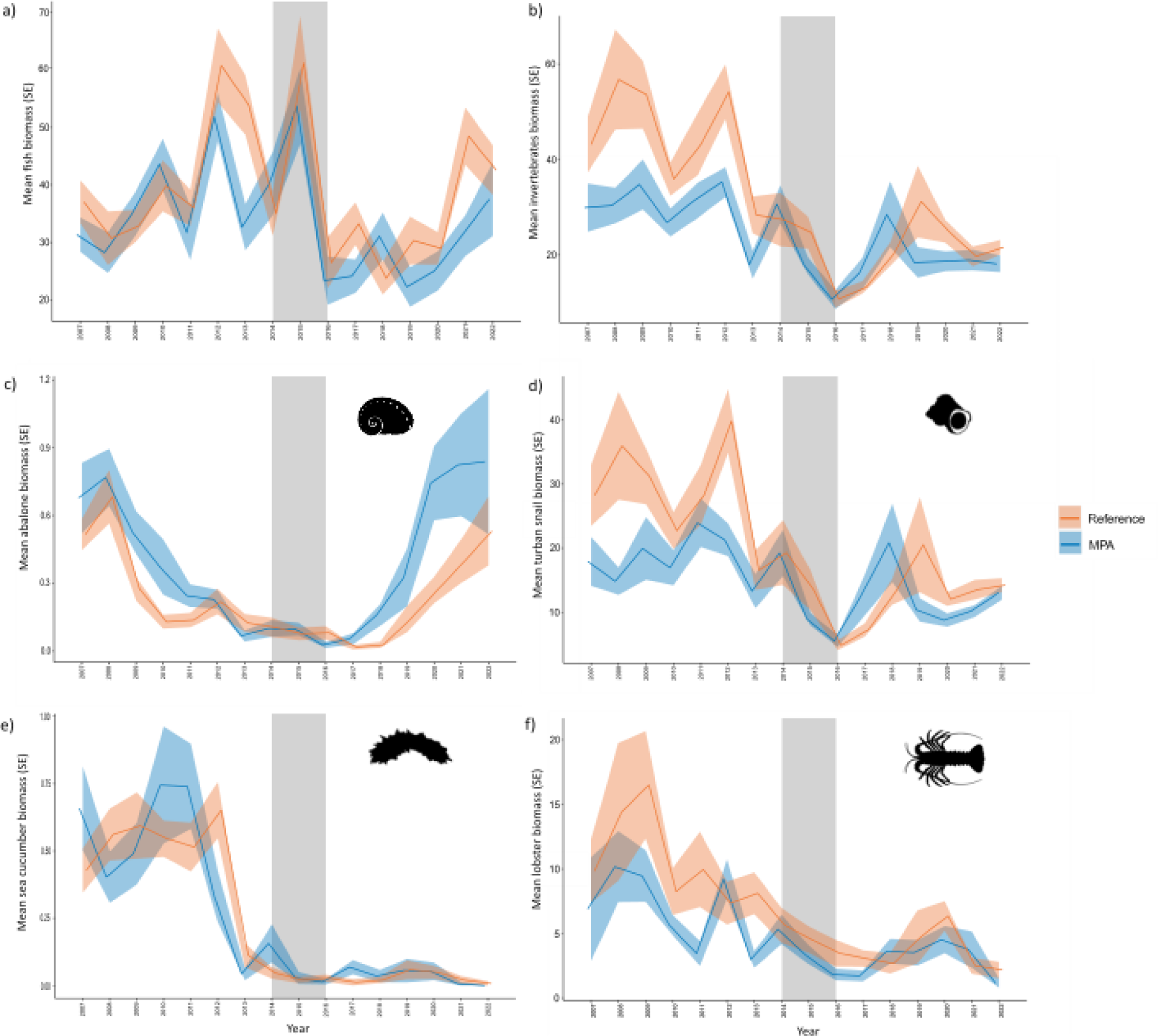
Time series (2007-2022) of biomass (mean biomass in kg in 60 m^2^, *n*= two reserves (blue) and two reference (orange) sites with 14-27 census transects each) for a) all fish species, b) all invertebrate species, c) abalone, d) turban snail, e) sea cucumber and f) California spiny lobster in Isla Natividad, Mexico. Shading shows standard error and nonoverlap indicates significant differences in mean biomass between sites for that year. The gray shaded area represents the 2014-2016 MHWs event.

**Figure 3.**
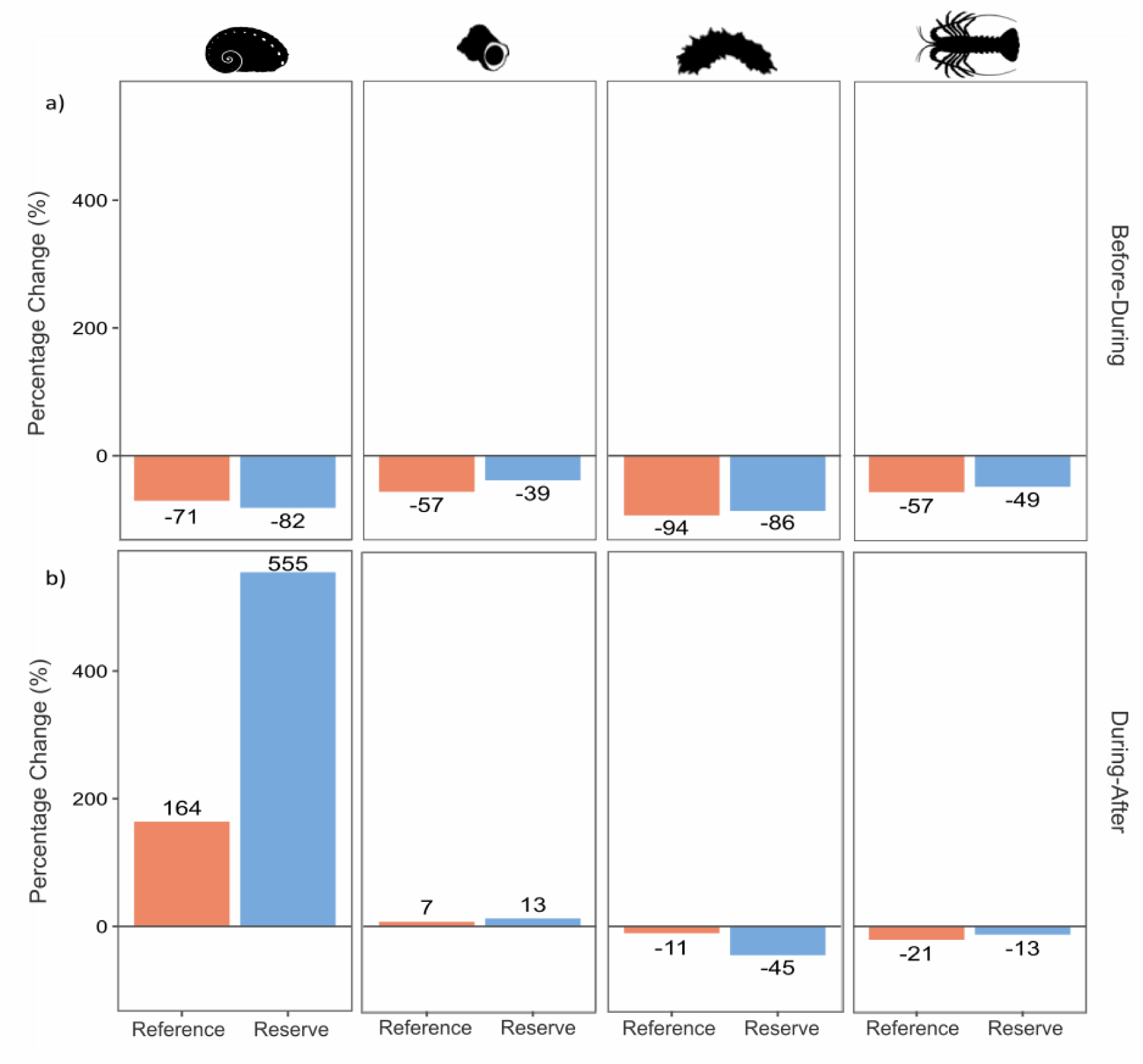
Percentage change in biomass a) before-during the MHWs and b) during-after the MHWs in reference sites (orange bars) and marine reserves (blue bars) for abalone, turban snail, sea cucumber and California spiny lobster in Isla Natividad, Mexico.

When exploring the impact of the MHWs on invertebrate species of interest, the GLM (Table 1) detected a negative impact on the turban snail, sea cucumber and lobster. The abalone showed a slight positive correlation during the MHWs, which may be related to biomass stabilization from 2013 to 2016, following the decline that occurred during the hypoxia event of 2009 (Micheli *et al*. 2012) and before the 2014-2016 MHW. Both the reference and reserves sites had a significant effect during the MHWs for the abalone, turban snail and lobster (Table 1), meaning that the MHWs impact was not less severe inside reserves than outside.

**Table 1.** Most parsimonious GLM for species of interest before and during the MHWs (2007-2016). Significance codes indicate significance at α = 0.001 (***), α = 0.01 (**), α = 0.05 (*), and marginally significant results at α = 0.1 (.).

| ***Haliotis* spp.: Biomass (log) ~ (Zone \* Marine Heatwave Status) + Year -1**<br>AIC: -46.408; $R^2 = 0.842$; df = 20; $e_i \sim N(0, \sigma^2)$; Family = Gaussian | | | | |
| --- | --- | --- | --- | --- |
| Predictors | Estimate | Std. Error | P-value | Significant |
| Zone: Reference | 0.445 | 0.042 | 1.97E-8 | *** |
| Zone: Reserve | 0.498 | 0.042 | 4.35E-9 | *** |
| MHW Status: During | 0.137 | 0.613 | 0.040 | * |
| Year | -0.057 | 0.008 | 6.02E-06 | *** |
| Zone (Reserve): MHW Status (During) | -0.061 | 0.063 | 0.352 |  |
| ***Megastrea* spp.: Biomass (log) ~ Zone + Marine Heatwave Status -1**<br>AIC: 18.431; $R^2 = 0.501$; df = 20; $e_i \sim N(0, \sigma^2)$; Family = Gaussian | | | | |
| Predictors | Estimate | Std. Error | P-value | Significant |
| Zone: Reference | 2.793 | 0.119 | 2.08E-14 | *** |
| Zone: Reserve | 2.554 | 0.119 | 9.07E-14 | *** |
| MHW Status: During | -0.636 | 0.166 | 0.0014 | ** |
| ***Apostichopus parvimensis*: Biomass (log) ~ (Zone \* Marine Heatwave Status) + Year -1**<br>AIC: NA, $R^2$: NA, Residual deviance: 0.922; df = 20; Family = Quasibibomial | | | | |
| Predictors | Estimate | Std. Error | P-value | Significant |
| Zone: Reference | -0.127 | 0.328 | 0.704 |  |
| Zone: Reserve | -0.335 | 0.331 | 0.326 |  |
| MHW Status: During | -2.118 | 0.937 | 0.039 | * |
| Year | -0.149 | 0.070 | 0.050 | . |
| Zone (Reserve): MHW Status (During) | 0.756 | 1.104 | 0.504 |  |
| ***Panulirus interruptus*: Biomass (log) ~ (Zone + Marine Heatwave Status) + Year -1**<br>AIC: 12.626; $R^2 = 0.507$; df = 20; $e_i \sim N(0, \sigma^2)$; Family = Gaussian | | | | |
| Predictors | Estimate | Std. Error | P-value | Significant |
| Zone: Reference | 1.609 | 0.099 | 8.76E-12 | *** |
| Zone: Reserve | 1.359 | 0.099 | 1.27E-10 | *** |
| MHW Status: During | -0.445 | 0.139 | 0.005 | ** |

### 3.4 Recovery of commercial species inside marine reserves and reference sites

The biomass of abalone and turban snails after the MHWs compared to during period as higher inside the reserves than in the reference sites. Exceptionally, abalone biomass had a ∼3.4-times higher increase of biomass after the MHWs inside the reserve (555%) compared to the reference sites (164%) (Fig 3b). On the other hand, the biomass of sea cucumber and lobster after the MHWs decreased compared to the during period. Sea cucumber was the only species that had less biomass inside the reserve than the reference site (Fig. 3b).

For the recovery period after the MHWs, three species (abalone, turban snail and lobster) reached the recovery threshold, showing signals of early recovery, inside the marine reserves (Fig. 4a, 4b, 4d). Abalone recovered after two years inside the reserves, and the recovery was sustained in time and increased every year until the end of this study. By 2021 and 2022, the abalone biomass recovery was outstanding, with a 10-fold increase compared to the biomass during the MHWs. In the reference sites, abalone also recovered but required three years to reach the recovery threshold and the recovery was less pronounced (5-fold) than in the reserve sites. Abalone is the only species that recovered both inside and outside the marine reserves and recovery was sustained through time (Fig. 4a). Turban snails recovered after two years, while lobsters after four years inside reserves (Fig. 4b & 4d). However, this recovery was not sustained through time. On the other side, the sea cucumber collapsed both in reserves and reference sites and never recovered to pre-MHW biomass (Fig. 4c). Importantly, only the abalone reached and surpassed the average pre-MHW baseline levels (2007-2013) after the MHWs (2017-2022), suggesting that turban snail and lobster populations have not fully recovered during the time frame of this study (Fig. S1).

**Figure 4.**
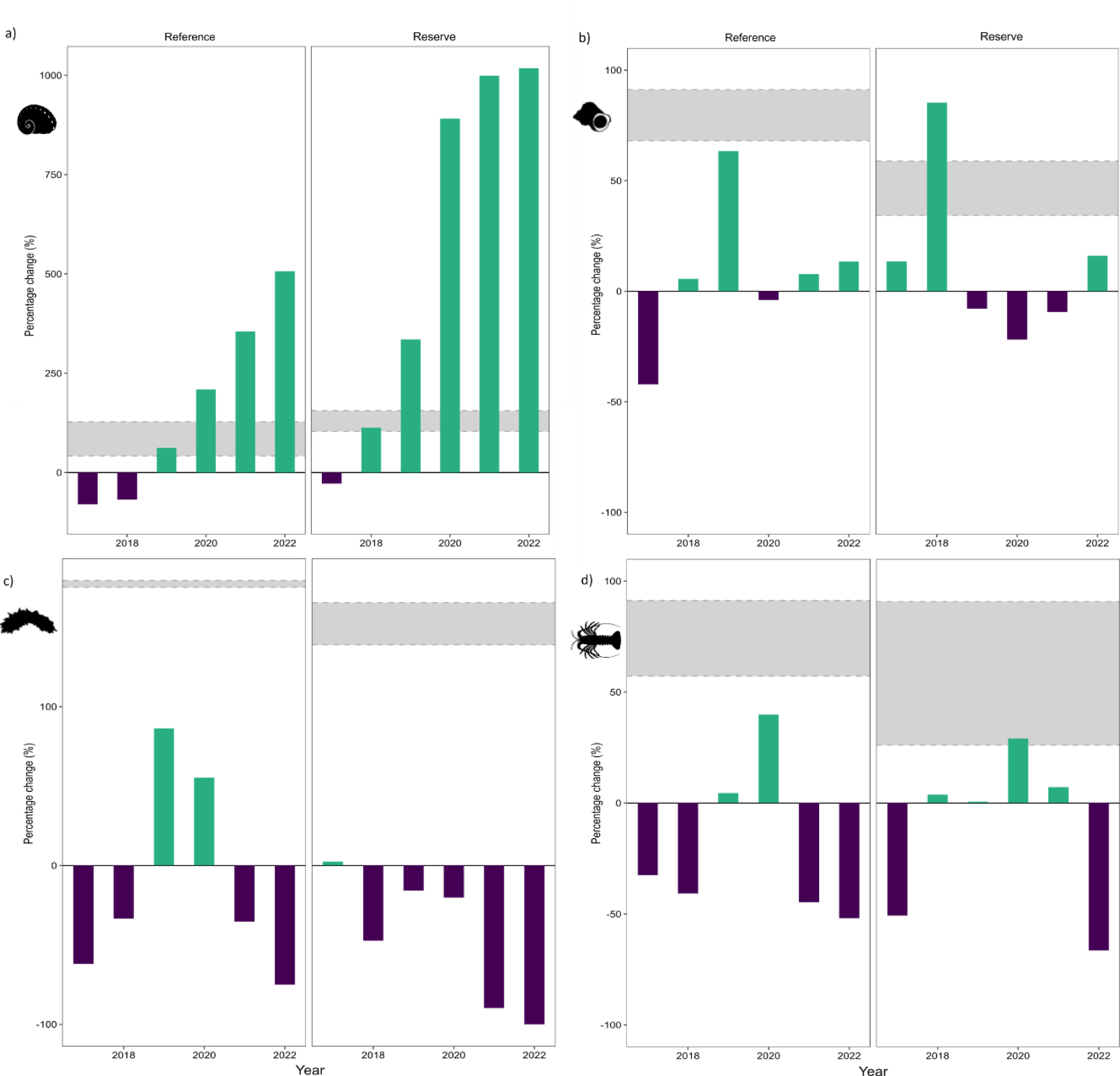
Percentage change (%) in annual mean biomass per transect after the MHWs (2017-2022) compared to mean values during (2014-2016) the MHWs of sedentary species a) abalone, b) turban snail, c) warty sea cucumber and mobile species d) California lobster in reference (left panel) and reserve (right panel) areas in Isla Natividad, Mexico. Positive (green) and negative (purple) bars represent years where abundance is above or below mean values during the heatwaves. Shading represents annual mean abundances with the 25^th^ and 75^th^ percentiles for the pre-MHWs (2007-2013) period.

## 4. Discussion

In our study, we showed that fully protected MPAs (i.e., marine reserves) can enhance the recovery of some kelp-associated species from MHWs impacts. Therefore, our findings support the notion that well managed fully protected MPAs can confer resilience to climate change impacts (Micheli *et al*. 2012; Roberts *et al*. 2017; Eisaguirre *et al*. 2020; Ziegler *et al*. 2023), and that MPAs can be effective climate adaptation tools (Arafeh-Dalmau *et al*. 2023a; CBD 2023). However, the impacts of and recovery from MHWs varied among species, suggesting that benefits are variable. This has important implications for MPA outcomes and expectations under a changing climate. In particular, our results suggest that MPAs may be more effective for conferring resilience to climate impacts to sedentary benthic species than mobile species that have larger home ranges. Our findings support global efforts of protecting 30% of the oceans by 2030 while adapting to climate change (CBD 2022) and contribute to the growing need to increase understanding the conditions for which MPAs can confer resilience to climate change impacts.

With future projections of increasing extreme climatic events such as MHWs (Oliver *et al*. 2019; IPCC 2023), understanding the benefits of MPAs for ecologically, economically and culturally important species under these threats is crucial. While several studies have documented the impacts of different MHWs globally (Smith *et al*. 2023b), only a few have explored whether MPAs confer resilience to MHW impacts (Freedman *et al*. 2020; Smith *et al*. 2023a; Benedetti-Cecchi *et al*. 2024; Kumagai *et al*. 2024). Here, we observed a significant decrease in the biomass of the invertebrate community, while the fish biomass was not impacted during the MHWs, partly supporting our hypothesis that species were impacted during the MHWs. These findings align with similar observations that reported higher vulnerability to MHW impacts of sedentary species which have limited capacity to escape from extreme events (Arafeh-Dalmau *et al*. 2019; Garrabou *et al*. 2022; Arafeh-Dalmau *et al*. 2023a).

Evidence for MPA effectiveness in providing climate resilience is conflicting but emerging. While Ziegler *et al*. (2023) reported faster recovery of fish species diversity inside MPAs compared to reference sites following the 2014-2016 MHWs in Central California, USA, other studies in California found that MPAs did not confer community structure resilience to the same MHWs for kelp forests, rocky reefs, and deeper rocky reefs (Freedman *et al*. 2020; Smith *et al*. 2023a), except for rocky intertidal habitats (Smith *et al*. 2023a). On the other hand, MPAs in California did provide resilience for kelp forests by preserving trophic cascades and preventing sea urchins from overgrazing kelp forests in southern California (Kumagai *et al*. 2024). In this study, we provide new evidence of population recovery inside MPAs, following MHWs, thereby contributing to the emerging science evaluating the efficacy of MPAs as climate adaptation tools.

The unprecedented 2014-2016 MHWs represents a unique ‘natural experiment’ that, combined with the existing long-term monitoring of the kelp forest ecosystem of Isla Natividad, provided an opportunity to explore the role of MPAs during and after MHWs. We detected faster signs of recovery from the MHWs for three invertebrate species within reserves compared to reference sites, supporting our hypothesis that reserves enhance species recovery. Abalone was the only species that recovered from MHWs in both reference and reserves sites and the recovery was sustained and increased through time. However, abalone recovery inside marine reserves outperformed by 3.3 times higher biomass than in reference sites. Moreover, the small marine reserves in Isla Natividad have supported the recovery of depleted abalone populations through increased biomass, body size, and reproductive output of two abalone species (Micheli *et al*. 2012; Smith *et al*. 2022). Micheli *et al*. (2012) found that these mechanisms also provided resilience to mass mortality from extreme hypoxia for local abalone populations by supporting a higher abundance of larger reproductive individuals inside the reserves and increased juvenile recruitment inside and around the reserves. The massive abalone biomass recovery reported here after the MHWs is likely attributed to the recovery of larger adults, a mechanism that may be providing climate resilience for other invertebrate species. This is of great importance, especially considering the marine reserves in Isla Natividad were primarily established for invertebrates, particularly abalone, rather than fish (Micheli *et al*. 2012). Therefore, these reserves are meeting their objective, even under multiple climate change impacts (i.e., hypoxia, MHWs).

It is essential to conduct extensive, long-term monitoring across ecosystems, multiple taxa, and regions to understand population dynamics (White 2018), and to evaluate the capacity of MPAs to promote recovery from climate extremes. The long-term monitoring of Isla Natividad is an example of the value of continuous monitoring for detecting trends in species abundance and biomass inside and outside MPAs over a 16-year period that included multiple MHWs. Similarly, long-term monitoring of the Leigh Marine Reserve (established in 1977 in New Zealand) showed that long term protection resulted in a stable kelp forests ecosystem, whereas fished sites exhibited prevalence of sea urchin barrens and algal turfs (Peleg *et al*. 2023). Long-term monitoring provides the data needed to evaluate the effectiveness of MPAs under a changing climate and to clarify expectations on the effectiveness of MPAs.

Our study demonstrated that MPAs in Baja California Sur, Mexico, did not mitigate the impact of MHWs, which did not support our hypothesis that the MHWs impact would be less severe inside reserves, but enhanced the recovery of three commercially important species following MHWs. Specifically, the recovery of abalone is outstanding empirical proof of the efficacy of MPAs. The findings that two small MPAs in Isla Natividad enhanced the resilience of invertebrate species to MHW impacts are encouraging and raise questions about the potential multiplicative benefits that additional MPAs (Arafeh-Dalmau *et al*. 2023a) could deliver, particularly if established in climate refugia areas (Woodson *et al*. 2019).

Furthermore, the observed differences in recovery rates among species in MPAs highlight the importance of designing and managing networks of climate-smart MPAs that adequately protect multiple species, are well-connected, and that include climate adaptation strategies (Arafeh-Dalmau *et al*. 2023a). Our study provides new insights regarding MPAs’ role in supporting climate resilience, that nations can use to achieve the protection of 30% of their oceans by 2030 (CBD 2023) while adapting to climate change.

## Acknowledgments

C.O-J., N.A.-D., A.H-V., A.R., J.T., C.B.W. and F.M. acknowledge the support of the US National Science Foundation (DISES 2108566). We are thankful to the members of Cooperativa de Producción Pesquera Buzos y Pescadores de la Baja California for their support and participation.

## Author Contribution

Conceptualization: C.O-J., N.A-D., F.M. Data Curation: C.O-J, M-M.E-C., A.H-V, A.R., F.M. Formal Analysis: M-M.E-C. Funding Acquisition: F.M. Investigation: C.O-J, M-M.E-C., A.H-V, A.R, F.M. Project Administration: C.O-J, F.M.. Writing first draft: C.O-J, N.A-D. Writing-review & editing: C.O-J, N.A-D, M-M.E-C., J.A.E.M., A.H-V, R.M., A.R., J.T., C.B.W., F.M.

## Statement of Inclusion

Our study brings together authors from two different countries, including scientists based in the country where the study was carried out. Data collection of this study took place in Isla Natividad, Mexico and we include local people as co-authors because the study would not exist without them. All authors were engaged early on with the research and study design to ensure that the diverse sets of perspectives they represent were considered from the onset. Whenever relevant, literature published by scientists from the region was cited. Authors in this study are part of the Fishing Cooperative *Buzos y Pescadores de la Baja California* where the study was carried out.

