## Supplementary Material for "Recovery mode: Marine protected areas enhance the resilience of kelp species from marine heatwaves"

**Supplementary Materials**

**Table S1.** Mass of fish and invertebrates used for biomass recovery based on Woodson *et al.* (2019) and Fajardo-León *et al.* (2008). Species in bold were used for the recovery analysis of this study.

| Fish Species | Mass (kg) | Invertebrate Species | Mass (kg) |
| --- | --- | --- | --- |
| *Anisotremus davidsoni* | 0.50 | Anemone *spp.* | 0.25 |
| *Caulolatilus princeps* | 3.0 | ***Apostichopus parvimensis*** | **0.25** |
| *Chromis punctipinnis* | 0.10 | *Cancer spp.* | 0.50 |
| *Embiotica jacksoni* | 0.20 | *Centrostephanus coronatus* | 0.40 |
| *Girella nigricans* | 1.50 | *Crassedoma giganteum* | 0.25 |
| *Halichoeres semicinctus* | 0.10 | *Cypraea spp.* | 0.25 |
| *Heterodontus francisci* | 4.00 | ***Haliotis spp.*** | **0.25** |
| *Hypsypops rubicundus* | 0.30 | *Kelletia kelletii* | 0.25 |
| *Mycteroperca spp.* | 3.00 | *Leptogorgia chilensis* | 0.25 |
| *Ophiodon elongatus* | 3.50 | *Loxorhyncus grandis* | 1.00 |
| *Oxyjulis californica* | 0.05 | ***Megastraea turbanica*** | **0.50** |
| *Paralabrax spp.* | 0.80 | ***Megastraea undosum*** | **0.30** |
| *Rhacochilus vacca* | 0.10 | *Megathura crenulate* | 0.30 |
| *Rhinobatos productus* | 8.00 | *Mesocentrotus franciscanus* | 0.25 |
| *Scorpaenichthys marmoratus* | 3.50 | *Muricea spp.* | 0.25 |
| *Sebastes spp.* | 1.00 | *Neobernaya spadicea* | 0.10 |
| *Semicossyphus pulcher* | 3.50 | *Octopus spp.* | 1.00 |
| *Squatina californica* | 5.00 | ***Panulirus interruptus*** | **1.00** |
| *Stereolepis gigas* | 8.00 | *Patiria miniata* | 0.40 |
|  |  | *Pisaster giganteus* | 1.00 |
|  |  | *Pycnopodia helianthoides* | 1.00 |
|  |  | *Strongylocentrotus purpuratus* | 0.40 |

**Table S2.** Start, peak and end date, duration and intensity (maximum, mean and accumulated) of MHWs events registered at Isla Natividad, Baja California Sur, Mexico from 1983-2022.

| **Start Date** | **Peak Date** | **End Date** | **Duration (days)** | **Maximum Intensity (°C)** | **Mean Intensity (°C)** | **Cumulative Intensity (°C)** |
| --- | --- | --- | --- | --- | --- | --- |
| 14-01-83 | 14-01-83 | 22-01-83 | 9 | 3.1513 | 2.4806 | 22.3252 |
| 26-01-83 | 29-01-83 | 08-03-83 | 42 | 2.4066 | 1.9114 | 80.2776 |
| 22-03-83 | 23-03-83 | 28-03-83 | 7 | 1.724 | 1.6159 | 11.3115 |
| 02-04-83 | 03-04-83 | 08-04-83 | 7 | 1.79 | 1.4523 | 10.1659 |
| 18-04-83 | 23-04-83 | 25-04-83 | 8 | 1.774 | 1.4725 | 11.7798 |
| 30-04-83 | 01-05-83 | 04-05-83 | 5 | 1.5023 | 1.3273 | 6.6366 |
| 08-08-83 | 23-08-83 | 01-09-83 | 25 | 3.848 | 3.0398 | 75.9945 |
| 09-09-83 | 12-09-83 | 10-10-83 | 32 | 4.4461 | 3.6016 | 115.2524 |
| 26-10-83 | 29-10-83 | 31-10-83 | 6 | 2.6535 | 2.545 | 15.27 |
| 10-11-83 | 12-11-83 | 15-11-83 | 6 | 2.7014 | 2.2432 | 13.4593 |
| 07-07-84 | 07-07-84 | 22-07-84 | 16 | 3.322 | 2.378 | 38.0477 |
| 18-08-84 | 23-08-84 | 28-08-84 | 11 | 3.288 | 2.7409 | 30.1501 |
| 02-09-84 | 07-09-84 | 08-09-84 | 7 | 3.7636 | 3.0749 | 21.5243 |
| 10-05-87 | 13-05-87 | 16-05-87 | 7 | 1.8513 | 1.6467 | 11.5267 |
| 05-04-88 | 08-04-88 | 16-04-88 | 12 | 1.4514 | 1.2832 | 15.3989 |
| 23-12-89 | 26-12-89 | 27-12-89 | 5 | 2.4427 | 2.1454 | 10.7269 |
| 10-07-90 | 10-07-90 | 19-07-90 | 10 | 2.7462 | 2.2038 | 22.0384 |
| 26-07-90 | 29-07-90 | 31-07-90 | 6 | 2.8021 | 2.4359 | 14.6156 |
| 06-12-90 | 09-12-90 | 15-12-90 | 10 | 2.073 | 1.765 | 17.6496 |
| 04-02-92 | 10-02-92 | 11-02-92 | 8 | 1.752 | 1.5851 | 12.6812 |
| 24-02-92 | 10-06-92 | 17-06-92 | 115 | 3.6323 | 2.3643 | 271.8931 |
| 15-07-92 | 20-07-92 | 23-07-92 | 9 | 2.5098 | 2.1662 | 19.496 |
| 22-09-92 | 29-09-92 | 02-10-92 | 11 | 3.6755 | 3.4034 | 37.4373 |
| 15-11-92 | 17-11-92 | 30-11-92 | 16 | 3.0556 | 2.3746 | 37.9942 |
| 06-12-92 | 10-12-92 | 13-12-92 | 8 | 2.1103 | 1.9354 | 15.4834 |
| 01-02-93 | 06-02-93 | 08-02-93 | 8 | 1.8279 | 1.7219 | 13.7755 |
| 14-01-94 | 16-01-94 | 24-01-94 | 11 | 2.0903 | 1.7869 | 19.6555 |
| 16-03-94 | 19-03-94 | 23-03-94 | 8 | 1.8151 | 1.594 | 12.7517 |
| 08-03-95 | 16-03-95 | 20-03-95 | 13 | 2.3154 | 1.908 | 24.8046 |
| 03-05-96 | 06-05-96 | 09-05-96 | 7 | 1.5855 | 1.47 | 10.2903 |
| 15-05-97 | 16-05-97 | 23-05-97 | 9 | 1.877 | 1.665 | 14.9852 |
| 27-05-97 | 01-06-97 | 05-06-97 | 10 | 1.7667 | 1.5483 | 15.4831 |
| 02-08-97 | 19-09-97 | 19-09-97 | 49 | 4.4057 | 3.164 | 155.0377 |
| 25-09-97 | 18-10-97 | 04-03-98 | 161 | 4.6094 | 2.8666 | 461.5255 |
| 08-03-98 | 11-03-98 | 12-03-98 | 5 | 2.2039 | 1.6947 | 8.4734 |
| 03-05-98 | 05-05-98 | 09-05-98 | 7 | 1.8371 | 1.4843 | 10.3903 |
| 17-05-98 | 18-05-98 | 22-05-98 | 6 | 1.5595 | 1.4388 | 8.6328 |
| 16-07-98 | 16-07-98 | 21-07-98 | 6 | 2.7268 | 2.3583 | 14.1498 |
| 12-08-98 | 12-08-98 | 16-08-98 | 5 | 2.8017 | 2.5952 | 12.9759 |
| 23-06-00 | 25-06-00 | 01-07-00 | 9 | 3.0305 | 2.299 | 20.6913 |
| 22-06-01 | 05-07-01 | 09-07-01 | 18 | 3.0939 | 1.9572 | 35.2298 |
| 12-02-03 | 17-02-03 | 19-02-03 | 8 | 2.1019 | 1.7867 | 14.2933 |
| 30-05-05 | 02-06-05 | 03-06-05 | 5 | 1.5503 | 1.4022 | 7.0109 |
| 13-07-06 | 16-07-06 | 19-07-06 | 7 | 2.8268 | 2.2788 | 15.9517 |
| 20-12-06 | 27-12-06 | 05-01-07 | 17 | 2.3748 | 1.7994 | 30.5903 |
| 23-10-08 | 30-10-08 | 03-11-08 | 12 | 3.9087 | 3.0592 | 36.711 |
| 20-11-08 | 27-11-08 | 06-12-08 | 17 | 2.4113 | 2.06 | 35.0204 |
| 07-01-10 | 12-01-10 | 19-01-10 | 13 | 2.3026 | 1.7663 | 22.9622 |
| 01-09-12 | 04-10-12 | 01-11-12 | 62 | 5.9514 | 3.8276 | 237.3084 |
| 11-07-13 | 17-07-13 | 20-07-13 | 10 | 2.7661 | 2.3932 | 23.932 |
| 23-04-14 | 06-05-14 | 07-05-14 | 15 | 1.9855 | 1.3905 | 20.8575 |
| 15-05-14 | 17-05-14 | 06-06-14 | 23 | 2.4384 | 1.9337 | 44.4746 |
| 11-06-14 | 15-06-14 | 25-06-14 | 15 | 2.7377 | 2.104 | 31.5595 |
| 29-06-14 | 05-07-14 | 12-07-14 | 14 | 4.0539 | 2.7983 | 39.1767 |
| 28-07-14 | 03-08-14 | 04-08-14 | 8 | 2.8796 | 2.6423 | 21.1381 |
| 13-08-14 | 14-09-14 | 10-06-15 | 302 | 4.8915 | 2.6875 | 811.6191 |
| 17-06-15 | 21-06-15 | 03-07-15 | 17 | 2.5741 | 2.0283 | 34.4817 |
| 21-07-15 | 25-07-15 | 28-07-15 | 8 | 2.8482 | 2.4825 | 19.8596 |
| 04-08-15 | 05-08-15 | 10-08-15 | 7 | 3.0268 | 2.7561 | 19.2925 |
| 08-09-15 | 14-10-15 | 13-03-16 | 188 | 5.6984 | 3.4048 | 640.103 |
| 08-04-16 | 22-04-16 | 16-05-16 | 39 | 2.9014 | 1.8113 | 70.6411 |
| 29-05-16 | 02-06-16 | 05-06-16 | 8 | 2.4003 | 1.9608 | 15.6862 |
| 19-06-16 | 22-06-16 | 09-07-16 | 21 | 2.5476 | 1.9918 | 41.8275 |
| 22-07-16 | 28-07-16 | 02-08-16 | 12 | 2.5151 | 2.3018 | 27.6219 |
| 08-06-17 | 17-06-17 | 24-06-17 | 17 | 1.898 | 1.4643 | 24.8932 |
| 04-07-17 | 18-07-17 | 10-08-17 | 38 | 4.0164 | 2.7157 | 103.196 |
| 26-11-17 | 30-11-17 | 30-11-17 | 5 | 2.4417 | 2.1107 | 10.5536 |
| 30-12-17 | 08-01-18 | 19-01-18 | 21 | 2.6194 | 1.9472 | 40.8915 |
| 02-02-18 | 03-02-18 | 11-02-18 | 10 | 2.0448 | 1.7149 | 17.1492 |
| 06-08-18 | 11-08-18 | 26-08-18 | 21 | 3.5672 | 2.8772 | 60.4222 |
| 16-11-18 | 18-11-18 | 22-11-18 | 7 | 2.2987 | 2.1871 | 15.3095 |
| 22-12-18 | 22-12-18 | 27-12-18 | 6 | 1.8291 | 1.7243 | 10.346 |
| 15-01-19 | 18-01-19 | 21-01-19 | 7 | 1.6974 | 1.5881 | 11.117 |
| 29-01-19 | 01-02-19 | 02-02-19 | 5 | 1.9017 | 1.7561 | 8.7803 |
| 13-03-20 | 14-03-20 | 17-03-20 | 5 | 1.7264 | 1.6054 | 8.0272 |
| 06-05-20 | 10-05-20 | 13-05-20 | 8 | 1.4902 | 1.3944 | 11.1549 |
| 13-06-20 | 13-07-20 | 19-07-20 | 37 | 2.9846 | 2.1433 | 79.3023 |
| 14-10-20 | 14-10-20 | 24-10-20 | 11 | 2.6784 | 2.5392 | 27.9311 |
| 11-08-21 | 14-08-21 | 23-08-21 | 13 | 3.4407 | 2.9719 | 38.6352 |
| 08-04-22 | 11-04-22 | 12-04-22 | 5 | 1.2926 | 1.2076 | 6.0379 |
| 10-06-22 | 11-06-22 | 14-06-22 | 5 | 1.4438 | 1.335 | 6.675 |
| 21-06-22 | 28-06-22 | 01-07-22 | 11 | 2.8184 | 2.0866 | 22.953 |
| 24-12-22 | 28-12-22 | 31-12-22 | 8 | 1.5757 | 1.4456 | 11.565 |

**Table S3.** Yearly cumulative MHW intensities (°C days) registered in Isla Natividad, Baja California Sur, Mexico from 1983-2022.

|  | **Cumulative Intensity** | **Year** |
| --- | --- | --- |
| 1 | 362.4728 | 1983 |
| 2 | 89.7221 | 1984 |
| 3 | 0 | 1985 |
| 4 | 0 | 1986 |
| 5 | 11.5267 | 1987 |
| 6 | 15.3989 | 1988 |
| 7 | 10.7269 | 1989 |
| 8 | 54.3036 | 1990 |
| 9 | 0 | 1991 |
| 10 | 394.9852 | 1992 |
| 11 | 13.7755 | 1993 |
| 12 | 32.4072 | 1994 |
| 13 | 24.8046 | 1995 |
| 14 | 10.2903 | 1996 |
| 15 | 501.6926 | 1997 |
| 16 | 199.9611 | 1998 |
| 17 | 0 | 1999 |
| 18 | 20.6913 | 2000 |
| 19 | 35.2298 | 2001 |
| 20 | 0 | 2002 |
| 21 | 14.2933 | 2003 |
| 22 | 0 | 2004 |
| 23 | 7.0109 | 2005 |
| 24 | 37.9167 | 2006 |
| 25 | 8.6253 | 2007 |
| 26 | 71.7314 | 2008 |
| 27 | 0 | 2009 |
| 28 | 22.9622 | 2010 |
| 29 | 0 | 2011 |
| 30 | 237.3084 | 2012 |
| 31 | 23.932 | 2013 |
| 32 | 595.6583 | 2014 |
| 33 | 908.2755 | 2015 |
| 34 | 334.4052 | 2016 |
| 35 | 141.9023 | 2017 |
| 36 | 140.8589 | 2018 |
| 37 | 19.8973 | 2019 |
| 38 | 126.4155 | 2020 |
| 39 | 38.6352 | 2021 |
| 40 | 47.2309 | 2022 |

**Table S4.** Most parsimonious Generalized Linear Model (GLM) for the two functional groups (fish and invertebrates) before and during MHWs (2007-2016). Significance codes indicate significance at 𝛼 = 0.001 (***), 𝛼 = 0.01 (**), 𝛼 = 0.05 (*), and marginally significant results at 𝛼 = 0.1 (.).

| **Fish:** Biomass (log) ~ (Zone * Marine Heatwave Status) + Year -1  AIC: 21.984; *R^2^* =0.122; df = 20; *e_i_ ~ N* (0, σ ^2^); Family = Gaussian | | | | | | |
| --- | --- | --- | --- | --- | --- | --- |
| **Predictors** | **Estimate** | | **Std. Error** | | **P-value** | **Significant** |
| Zone: Reference | 3.344 | 0.230 | | 2.92E-10 | | *** |
| Zone: Reserve | 3.160 | 0.230 | | 6.48E-10 | | *** |
| MHW Status: During | -0.320 | 0.339 | | 0.359 | |  |
| Year | 0.038 | 0.046 | | 0.429 | |  |
| Zone (Reserve): MHW Status (During) | 0.065 | 0.350 | | 0.855 | |  |
| **Invertebrates:** Biomass (log) ~ Zone + Marine Heatwave Status -1  AIC: 15.639; *R^2^* = 0.555; df = 20; *e_i_ ~ N* (0, σ ^2^); Family = Gaussian | | | | | | |
| **Predictors** | **Estimate** | **Std. Error** | | **P-value** | | **Significant** |
| Zone: Reference | 3.406 | 0.111 | | 2.39E-16 | | *** |
| Zone: Reserve | 3.152 | 0.111 | | 8.74E-16 | | *** |
| MHW Status: During | -0.657 | 0.155 | | 0.0006 | | *** |

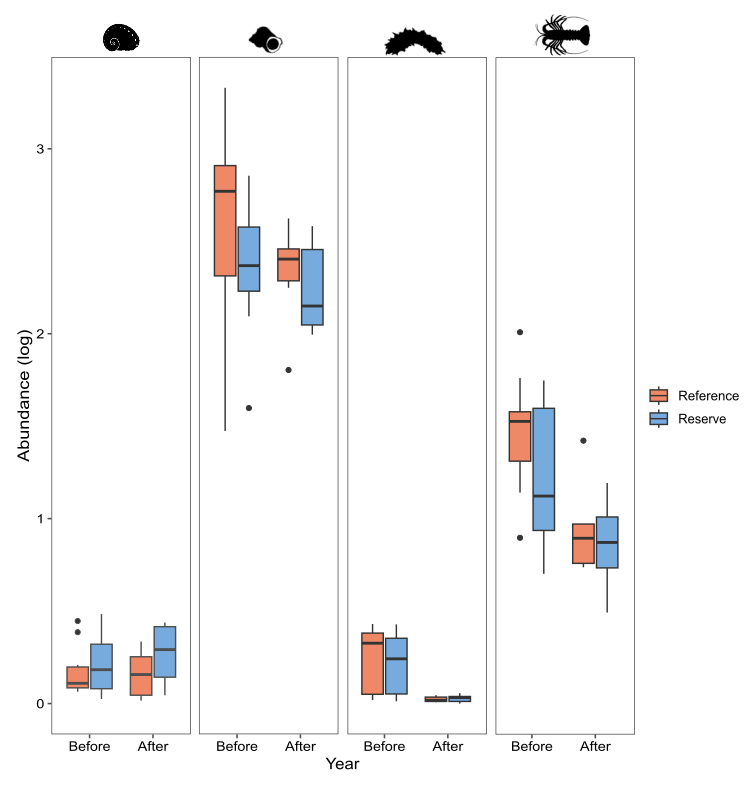

**Figure S1.** Boxplot showing the mean annual abundance before (2007-2013) and after (2017-2022) the MHWs inside marine reserves (blue bars) and reference sites (orange bars) for abalone, turban snail, warty sea cucumber and California spiny lobster in Isla Natividad, Mexico.
